# Manta: Rapid detection of structural variants and indels for clinical sequencing applications

**DOI:** 10.1101/024232

**Authors:** Xiaoyu Chen, Ole Schulz-Trieglaff, Richard Shaw, Bret Barnes, Felix Schlesinger, Anthony J. Cox, Semyon Kruglyak, Christopher T. Saunders

## Abstract

**Summary:** We describe Manta, a method to discover structural variants and indels from next generation sequencing data. Manta is optimized for rapid clinical analysis, calling structural variants, medium-sized indels and large insertions on standard compute hardware in less than a tenth of the time that comparable methods require to identify only subsets of these variant types: for example NA12878 at 50x genomic coverage is analyzed in less than 20 minutes. Manta can discover and score variants based on supporting paired and split-read evidence, with scoring models optimized for germline analysis of diploid individuals and somatic analysis of tumor-normal sample pairs. Call quality is similar to or better than comparable methods, as determined by pedigree consistency of germline calls and comparison of somatic calls to COSMIC database variants. Manta consistently assembles a higher fraction of its calls to basepair resolution, allowing for improved downstream annotation and analysis of clinical significance. We provide Manta as a community resource to facilitate practical and routine structural variant analysis in clinical and research sequencing scenarios.

**Availability:** Manta source code and Linux binaries are available from http://github.com/sequencing/manta.

**Supplementary information:** Supplementary data are available at *Bioinformatics* online.

## 1 INTRODUCTION

Whole genome and enrichment sequencing is increasingly used for discovery of inherited and somatic genome variation in clinical contexts, however tools for rapid discovery of structural variants (SVs) and indels in this scenario are limited. We address this gap with Manta, a novel method for accurate discovery and scoring of SVs, medium-sized indels and large insertions in a unified and rapid process. Manta discovers variants from a sequencing assay’s paired and split-read mapping information using an efficient parallel workflow. Many advanced structural variant methods are available which focus on research and population genomics (Rausch *et al.*, 2012; Layer *et al.*, 2014; Ye *et al.*, 2009; Sindi *et al.*, 2012).

However, none to our knowledge combine as many variant types into a rapid workflow focused on individual or small sets of related samples. Per its focus on clinical pipelines, Manta provides a complete solution for discovery, assembly and scoring using only a reference genome and alignments from any standard read mapper. It provides scoring models for germline analysis of diploid individuals and somatic analysis of tumor-normal sample pairs, with additional applications under development for RNA-Seq, *de novo* variants, and unmatched tumors. We describe Manta’s methods and compare with representative tools to demonstrate high variant call quality with dramatically reduced compute cost.

## 2 METHODS

### Workflow Summary

Manta’s workflow is designed for high parallelization on individual or small sets of samples. It operates in two phases: first a graph of all breakend associations within the genome is built, then the components of this graph are processed for variant hypothesis generation, assembly, scoring and VCF reporting. The breakend association graph contains edges between any genomic regions where evidence of a long range adjacency exists, indel assembly regions are denoted in this scheme as self-edges. The graph does not express specific variant hypotheses so it is very compact, and can be constructed from segments of the genome in parallel. Following graph construction, individual edges (or larger subgraphs) are analyzed for variants in parallel. Each edge is analyzed to find imprecise variant hypotheses, for which variant reads are assembled and aligned back to the genome. Assembly is attempted for all cases, but is not required to report a variant. All paired and split-read evidence is consolidated to a quality score under either a germline or somatic variant model, and filtration metrics complement this quality score to improve call precision. For ease of use, Manta automates estimation of insert size distribution and exclusion of high depth reference compression regions. Details of all workflow components are provided in Supplementary Methods.

### Variant Call Evaluation

We assess accuracy of germline calls by running variant callers on all members of CEPH pedigree 1463, selecting for calls with pedigree-consistent genotypes and evaluating each caller on one pedigree member (NA12878) against the pedigree-consistent call set. To find pedigree-consistent calls and provide a relative recall comparison for Manta, we select a standard recognized caller in each variant class: Pindel (Ye *et al.*, 2009) for indels and Delly (Rausch *et al.*, 2012) for SVs. Calls from each representative method are used to establish the pedigree-consistent call set together with Manta’s. For somatic calls we also use Delly as a standard benchmark and compare calls from both methods on breast cancer cell line HCC1954 to somatic variant entries for this sample in COSMIC v70 (Forbes *et al.*, 2015). Full details of the evaluation procedure are included in Supplementary Methods.

## 3 RESULTS

We describe NA12878 variant call performance in the top portion of Table 1, comparing the results of each method to pedigree-consistent calls for this sample (see Methods). The first section describes large deletions and duplications, showing that Manta’s results are competitive overall and have a somewhat higher recall (or higher rate of pedigree consistency due to correct genotyping). Manta calls consistently show a higher fraction of calls agreeing with the pedigree-consistent set which also have breakends assembled to basepair resolution. For deletions and insertions smaller than 500 bases, the next section of Table 1 reiterates the large SV pattern of strong performance, with a trend towards higher recall across these smaller indel variant classes.

**Table 1.**
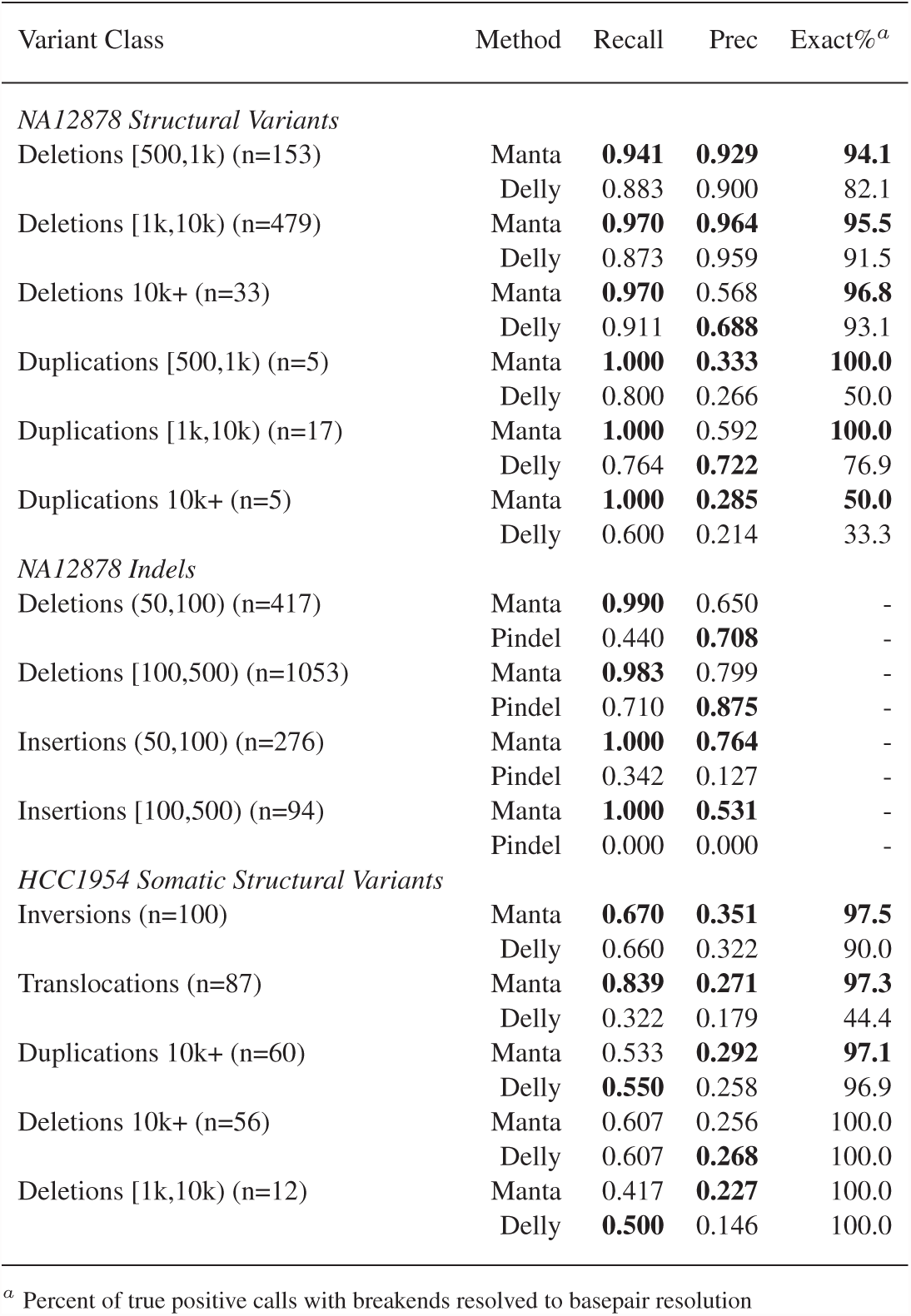
Assessment of variant call accuracy

Somatic call performance for HCC1954 is described in the final portion of Table 1, comparing each method’s variant calls to COSMIC variant entries for this cell line (see Methods). In this case, the truth set does not reflect a complete catalog of somatic variants for the cell line, however it does provide a useful relative precision estimate reflecting enrichment for known variants. Here we observe strong performance for Manta calls across all variant types with a trend towards a greater fraction of true calls assembled to basepair resolution, consistent with germline variant observations.

Table 2 summarizes runtime and memory cost for each variant caller, benchmarked in both parallel and serial modes to show workload distribution and methods efficiency. By either of these runtime or memory metrics we observe that Manta has substantially lower compute cost and turnaround time, while providing coverage of more variant types. We note that Delly is designed to parallelize primarily across, instead of within, samples, so the parallel test reflects a limited use of all server cores. When Manta is restricted to provide variant call coverage similar to Delly (variants 300 bases and larger), observed compute cost is even lower, further highlighting the efficiency of Manta’s implementation relative to current methods.

**Table 2.** Compute cost evaluation

| Sample | Method | Walltime (h) |  | Memory (Gb) |  |
| --- | --- | --- | --- | --- | --- |
|  |  | Parallel | Serial | Parallel | Serial |
| NA12878 | Manta | 0.327 | 3.764 | 2.351 | 0.233 |
|  | Manta-SV<sup>a</sup> | 0.102 | 0.878 | 1.786 | 0.125 |
|  | Pindel | 12.441 | 124.401 | 61.840 | 62.538 |
|  | Delly | 3.133 | 6.117 | 11.188 | 6.431 |
| HCC1954 | Manta | 0.852 | 5.486 | 3.445 | 0.244 |
|  | Manta-SV<sup>a</sup> | 0.544 | 2.391 | 2.754 | 0.186 |
|  | Delly | 75.911 | 100.648 | 11.614 | 8.540 |

All tests on dual Xeon E5-2680 v2 server with data on local drive. Parallel tests use all 20 cores, serial tests use 1 core. Memory columns show peak RSS.
<sup>a</sup> By default Manta assembles SVs and indels 8 bases and larger, Manta-SV is a custom SV-only configuration (300 bases and larger)

